# Independent Validation of Transgenerational Inheritance of Learned Pathogen Avoidance in *Caenorhabditis elegans*

**DOI:** 10.1101/2025.04.03.647070

**Authors:** A Akinosho, J Alexander, K Floyd, AG Vidal-Gadea

## Abstract

Transgenerational inheritance of learned behaviors remains a controversial topic in biology. The Murphy lab previously demonstrated that *C. elegans* exposed to pathogenic *Pseudomonas aeruginosa* (PA14) not only learn to avoid this bacterium but also transmit this avoidance behavior to untrained offspring. A recent study has challenged these findings, questioning the robustness of this phenomenon. Here, we independently validate the transgenerational inheritance of PA14 avoidance in *C. elegans*. Adapting the protocol described by the Murphy lab, we show that worms trained on PA14 develop significant avoidance that persists in F1 and F2 generations, though with decreased strength after P0. Our results provide independent confirmation of transgenerational inheritance of PA14 avoidance through the F2 generation, in agreement with previous findings from the Murphy lab and in contrast to a recent study by the Hunter group that failed to detect this phenomenon beyond the F1 generation.

## Introduction

*Caenorhabditis elegans* inhabits soil environments where it encounters diverse bacteria, including both nutritional and pathogenic species (Schulenburg and Félix, 2017; Neher, 2010). This nematode has evolved sophisticated sensory mechanisms to distinguish beneficial from harmful bacteria (Kim and Flavell, 2020; Guillermin, 2018). Interestingly, naïve *C. elegans* initially prefer the pathogenic bacterium *Pseudomonas aeruginosa* (PA14) over the non-pathogenic laboratory strain *Escherichia coli* (OP50) but rapidly learn to avoid PA14 following exposure (Zhang and Bargmann, 2005).

The Murphy lab first demonstrated that this learned avoidance behavior can be transmitted to progeny, persisting through the F2 generation (Moore et al. Cell 2019; Kaletsky et al. Nature 2020; Moore et al. Cell 2021; Sengupta et al., 2024). This transgenerational epigenetic inheritance (TEI) has been linked to small RNA pathways and the dsRNA transport proteins SID-1 and SID-2. Recently, Sengupta et al. (2024) further identified a specific small RNA responsible for inducing transgenerational inheritance of learned avoidance.

However, Gainey et al. (2025), representing the Hunter group, reported that while parental and F1 avoidance behaviors were evident, transgenerational inheritance was not reliably observed beyond the F1 generation under their experimental conditions. Critically, Kaletsky et al. (2025) demonstrated that omission of sodium azide during scoring can completely abolish detection of inherited avoidance, revealing that this key methodological difference may explain the conflicting results between laboratories. The use of sodium azide to immobilize worms at the moment of initial bacterial choice appears essential for capturing the inherited behavioral response. These findings highlight how seemingly minor methodological variations can dramatically impact detection of transgenerational inheritance and underscore the need for independent replication using standardized protocols. Here, we adapted the protocol established by the Murphy group, maintaining the critical use of sodium azide to paralyze worms at the time of choice, to test whether parental exposure to PA14 elicits consistent avoidance in subsequent generations. Our study specifically focuses on the transmission of learned avoidance through the F2 generation, beyond the intergenerational (F1) effect, because this is where divergence between published studies begins. Our goal was to evaluate whether the reported transgenerational signal persists across two generations without reinforcement.

## Results and Discussion

We investigated whether *C. elegans* exposed to PA14 exhibit learned avoidance behavior that persists across generations. Naïve worms showed a preference for PA14 over OP50, with a negative choice index (approximately -0.4). After 24 hours of exposure to PA14, trained worms developed strong aversion to the pathogen, exhibiting a significantly positive choice index (Figure 1).

**Figure 1.**
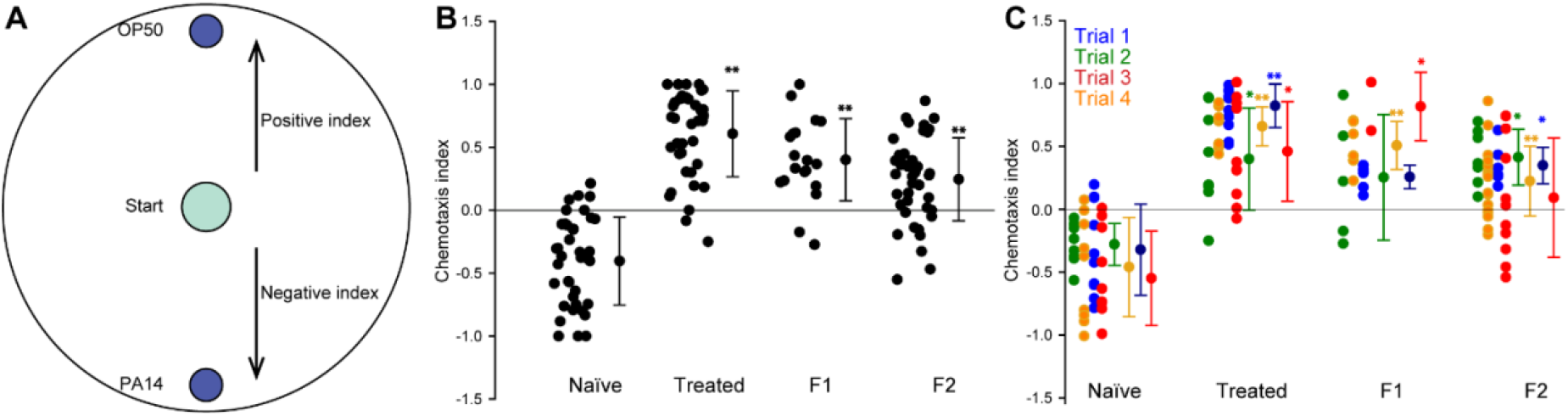
Transgenerational inheritance of PA14 avoidance in *C. elegans*. **A.** Chemotaxis assay setup. A 25 μL drop of OP50 and PA14 bacteria of the same optical density (OD600 = 1) plus 1.0 μL of 400 mM NaN3 were placed equidistant from the center of a 10cm chemotaxis plate where worms were placed at the start of the assay (see methods). indices were measured for naïve worms, trained worms, first-generation (F1), and second-generation (F2) offspring. **B**. Pooled results from four independent trials showing that Trained worms, which were exposed to PA14 for 24 hours, exhibited significantly higher choice indices, indicating learned avoidance of PA14. This aversion persisted in F1 and F2 progeny, demonstrating transgenerational inheritance of pathogen avoidance behavior. **C**. Comparison of data separated into individual trials. Kruskal-Wallis one-way ANOVA followed by Dunn’s post hoc, *p<0.05, **p<0.01 means and SEM shown (see Table S1 for statistics and Table S2 for raw data).

Critically, this avoidance behavior persisted in the untrained F1 and F2 progeny of trained worms, with both generations showing significantly higher choice indices compared to naïve controls (Fig. 1, Table S1 and S2).

Our findings independently validate the transgenerational inheritance of learned pathogen avoidance initially reported by the Murphy lab. In contrast, Gainey et al. (2025) using a modified protocol failed to observe such inheritance beyond F1 despite multiple attempts. While we cannot definitively reconcile these differences, our results suggest that under tightly controlled conditions, including bacterial lawn density, OD_600_=1.0 standardization, and immobilization with sodium azide restricted to one hour (to capture initial preference behaviors), transgenerational inheritance through F2 is both detectable and statistically robust.

This underscores the assay’s sensitivity to environmental variables, such as synchronization method and bacterial lawn density. This highlights the importance of consistency across experimental setups and support the view that context-dependent variation may underlie previously reported discrepancies.

These results confirm that, under tightly controlled conditions, transgenerational inheritance of PA14 avoidance extends through the F2 generation, though the response attenuates without further reinforcement.

## Methods

### Nematode Strains and Maintenance

Wild-type *Caenorhabditis elegans* (N2) were maintained on nematode growth medium (NGM) plates seeded with *Escherichia coli* OP50 at 20°C under standard conditions (Brenner 1974). Worm populations were passaged regularly to avoid overcrowding and starvation. OP50 was obtained from the Caenorhabditis Genetics Center (CGC).

### Bacterial Cultures

We inoculated 6 mL of LB medium (pH 7.5) with single colonies of OP50 or *Pseudomonas aeruginosa* PA14 and incubated them overnight at 37°C with shaking at 250 rpm. For PA14, cultures did not exceed 16 hours of incubation or develop visible biofilms. For both OP50 and PA14, the overnight culture was diluted to an OD_600_ of 1.0 in fresh LB media prior to seeding.

### Synchronization of Worms

Gravid adults were collected from OP50-seeded NGM plates by washing with liquid NGM buffer. We used liquid NGM buffer instead of M9 buffer (as specified in the original Murphy protocol) to maintain chemical consistency with the solid NGM culture plates. This modification minimizes potential osmotic stress since liquid NGM matches the pH (6.0) and ionic composition of the growth medium, whereas M9 buffer has a different pH (7.0) and ionic profile. After settling by gravity, a standard bleach solution was added and gently nutated for 5-10 minutes. The embryo pellet was washed 2-3 times with liquid NGM after centrifugation. Liquid NGM buffer is identical to agar NGM plates except that it does not include agar, peptone, cholesterol, or OP50 E. coli. The key difference between liquid NGM and M9 (aside from their pH) is that NGM has CaCl_2_ and that M9 typically has more Na+ while NGM has more K+ due to the phosphate salts used. Approximately 250-350 eggs were plated onto each OP50-seeded NGM plate and incubated at 20°C for 48-52 hours until reaching late L4 stage.

### Training and Choice Assay

Training plates (10 cm NGM) were seeded with 1 mL of overnight bacterial cultures and incubated at 25°C for 2 days. Choice assay plates (6 cm NGM) were prepared with 25 µL spots of OP50 and PA14 on opposite sides placed 24 hrs prior to the assays. Late L4 worms were transferred to training plates (OP50 or PA14) for 24 hours at 20°C. For choice assays, 1.0 µL of 400 mM sodium azide was placed on each bacterial spot before adding worms. After one hour, the number of worms immobilized on each spot was counted to calculate a Choice Index (CI):

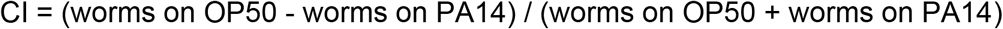

We used 400 mM sodium azide rather than the 1 M concentration reported by Moore et al. (2019) because preliminary trials showed that higher concentrations caused premature paralysis before worms could reach either bacterial spot, potentially biasing choice measurements. The 400 mM concentration provided sufficient immobilization while preserving the behavioral choice window.

### Transgenerational Testing

Synchronized F1 progeny were obtained by bleaching trained adult worms and allowing embryos to develop on OP50-seeded plates at 20°C. On Day 1 of adulthood, F1 worms were tested using the same choice assay. The F2 generation was derived from untrained F1 adults, synchronized in the same manner, and tested in identical conditions.

Each behavioral assay was conducted using animals from a biologically independent growth plate. While F2 plates were derived from pooled embryos from multiple F1 parents, each assay represents an independent biological replicate with no reuse of animals across assays. F2 assays (n=45) exceeded F1 assays (n=20) due to PA14-induced fecundity reduction in trained worms, limiting the number of viable F1 progeny. The higher number of F2 assays reflects the greater reproductive success of healthy F1 animals and provides additional statistical power for population-level behavioral comparisons.

### Controls and Additional Considerations

The data presented in this study are the result of four independent rounds of experimentations. Each individual assay was performed with animals harvested from unique culture plates, ensuring biological independence across all behavioral measurements. An average of 62 ± 43 animals participated in each assay (were immobilized in the NaN_3_ and tallied at the end of the assay). We conducted an average of 8.5 assays per condition during each of the four replicates. Each replicate was a biologically independent experiment conducted on a different day. Our experimental design employed population-level comparisons across generations using unpaired statistical analyses, with no attempt to track individual lineages across generations. All plates used in the study were prepared at least two days before experiments to ensure consistent bacterial lawn growth. Worms were monitored to prevent starvation or overcrowding, as these conditions could influence their bacterial preferences and behavioral responses. Fresh bacterial stocks were maintained at -80°C and streaked weekly to ensure culture consistency.

### Statistical Analysis

All statistical analyses were performed using SigmaPlot 14 (Inpixon). Outliers were identified and removed using the interquartile range criterion (>1.5×IQR), resulting in the exclusion of 5 out of 143 data points (3.5%). Their removal did not alter the significance or direction of reported effects. When datasets passed both the Shapiro-Wilk test for normality and the Brown-Forsythe test for equal variance, we used one-way ANOVA followed by Holm-Sidak post hoc comparisons versus the control group. For datasets that failed either assumption, we used Kruskal–Wallis one-way analysis of variance on ranks, followed by Dunn’s post hoc comparisons versus control. The full statistical outputs are provided in Table S1, and raw data for individual assays are presented in Table S2.

## Acknowledgments

We thank Dr. Murphy at Princeton University for reagents and protocols. Some strains were provided by the CGC, which is funded by NIH Office of Research Infrastructure Programs (P40 OD010440).

## Supplementary Information

**Table S1.**
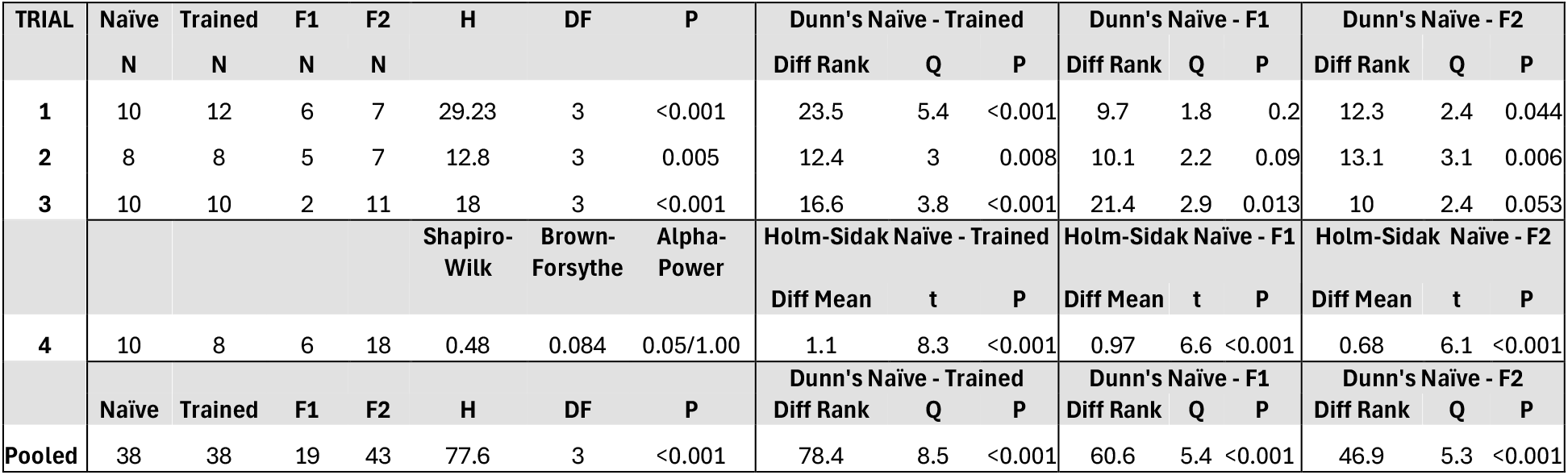
Summary of statistical analysis. H: H-statistic; DF: degrees of freedom; Q: standardized difference between ran sums; Trial: Independent trials conducted on different dates; N: independent assays conducted on a trial (each trial derived from individual culture plate); **Naïve**: Day-1 adults without previous exposure to PA14; **Trained**: Day-1 adults with previous exposure to PA14; **F1**: Day-1 adults without previous exposure to PA14 derived from **Trained** group; **F2**: Day-1 adults without previous exposure to PA14 derived from **F1** group.

**Table S2.**
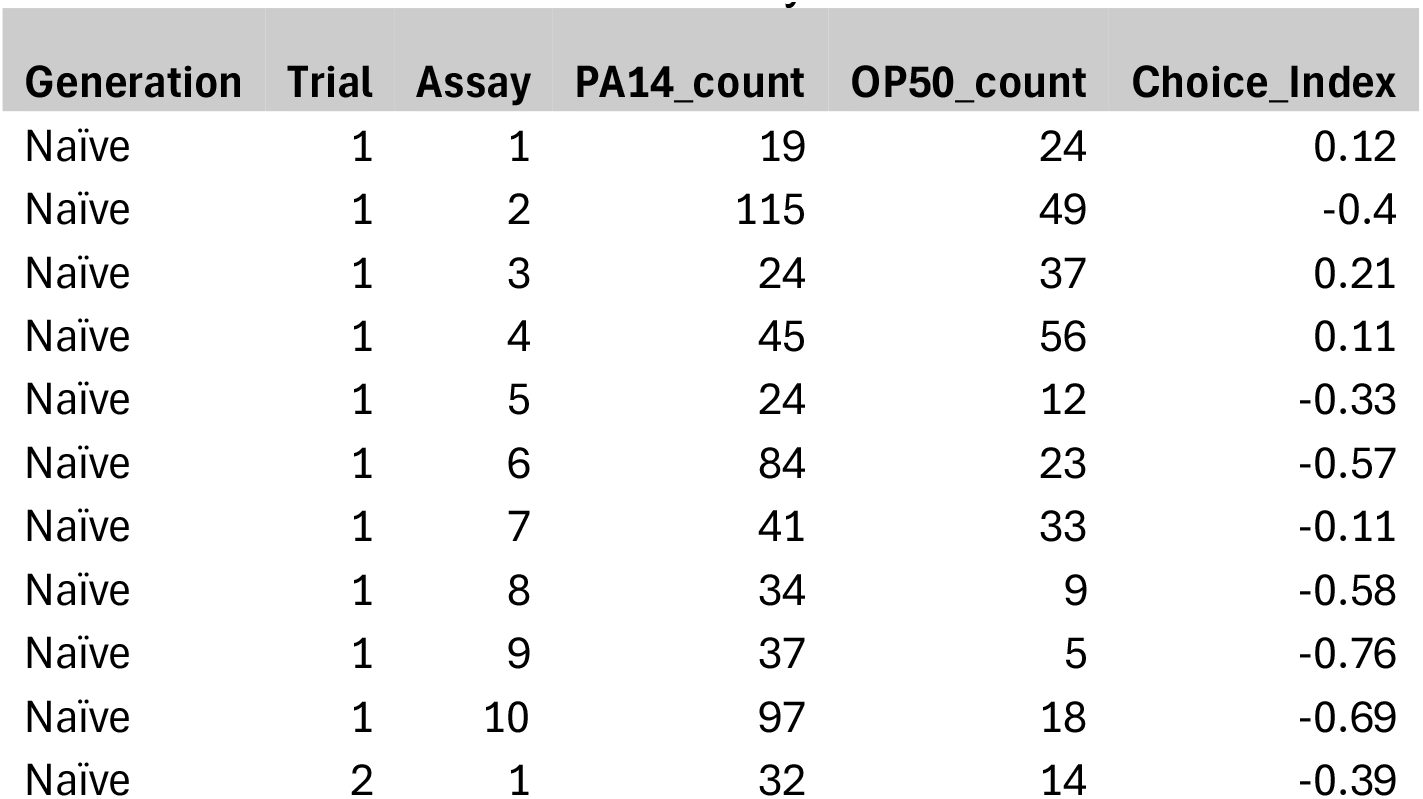
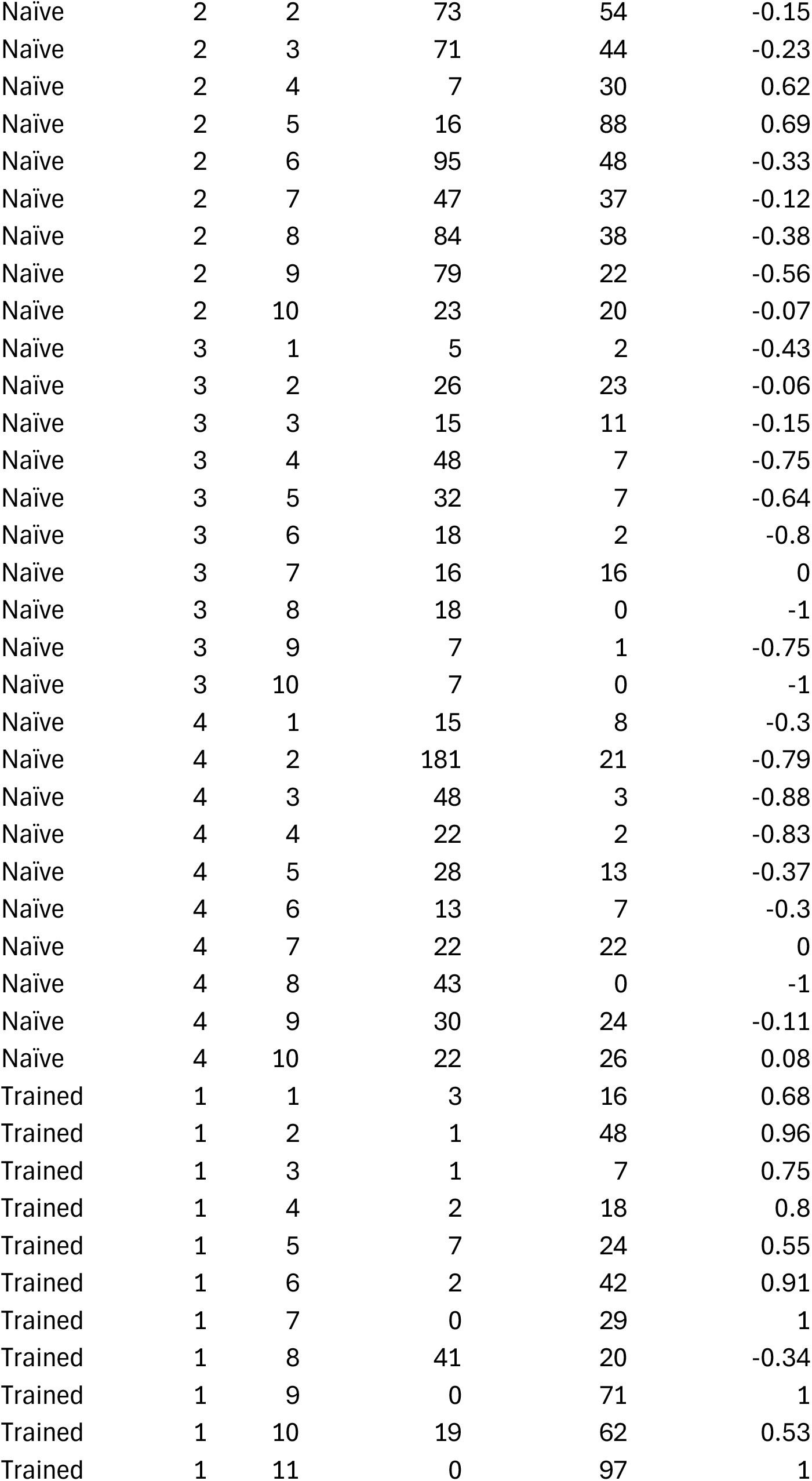
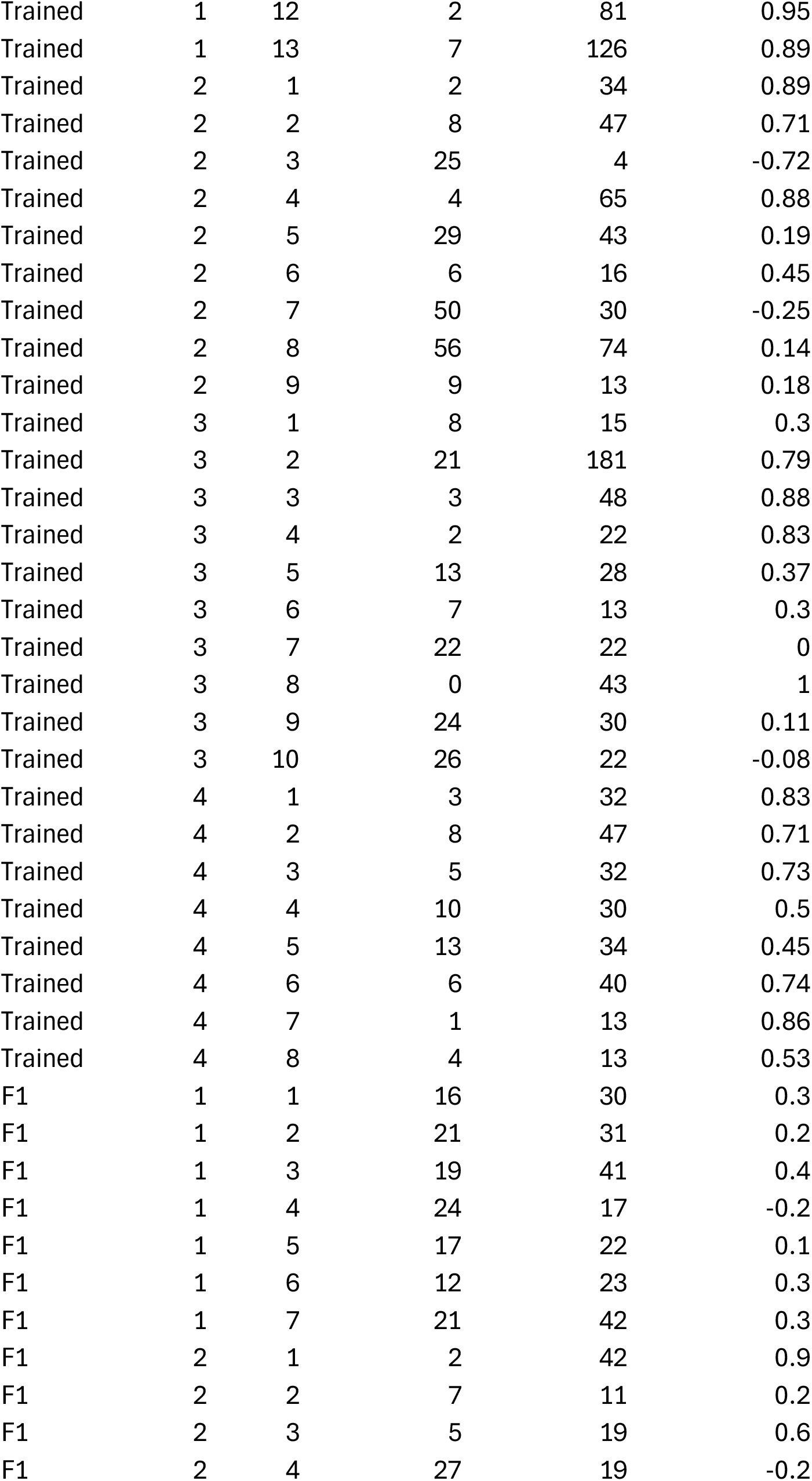
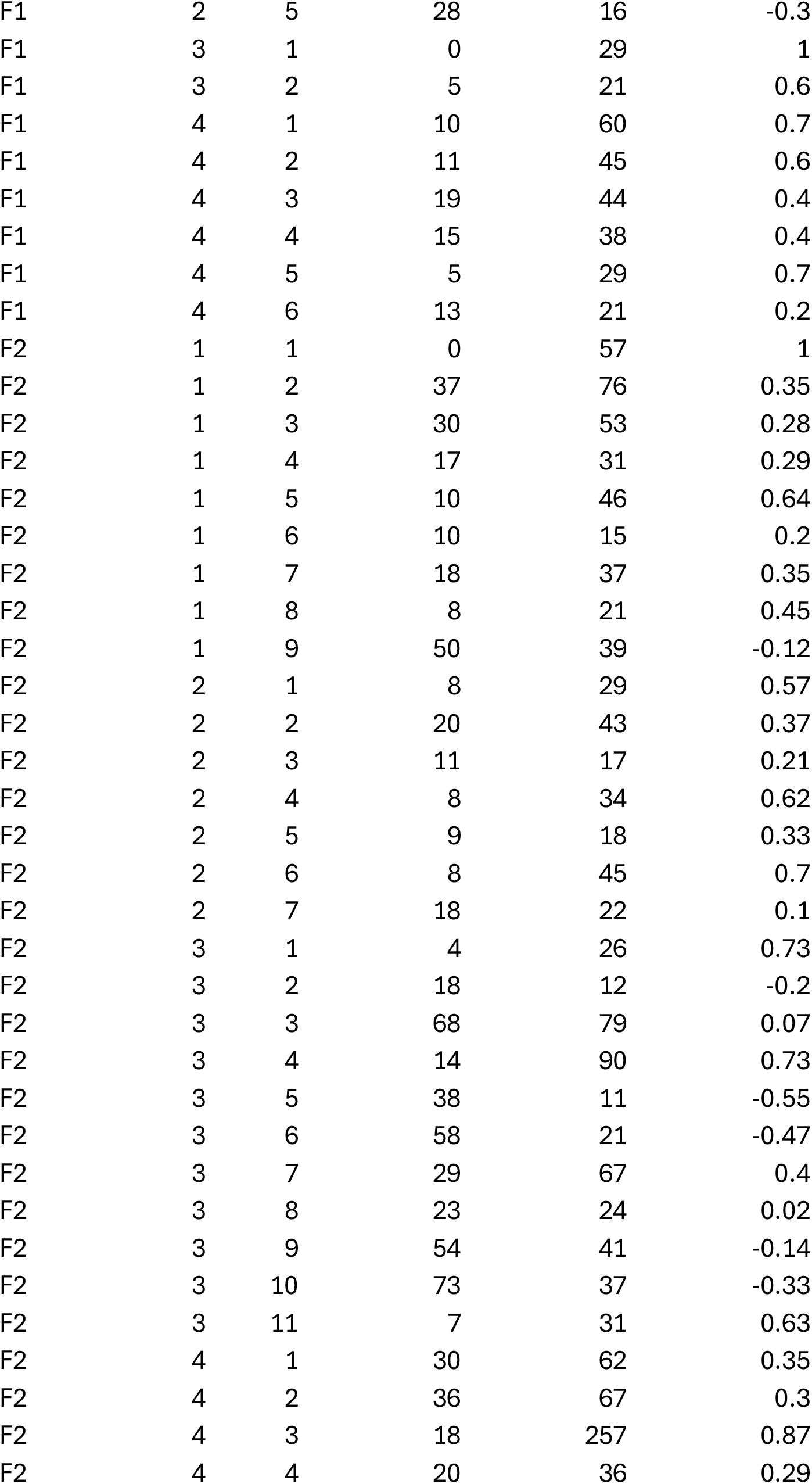
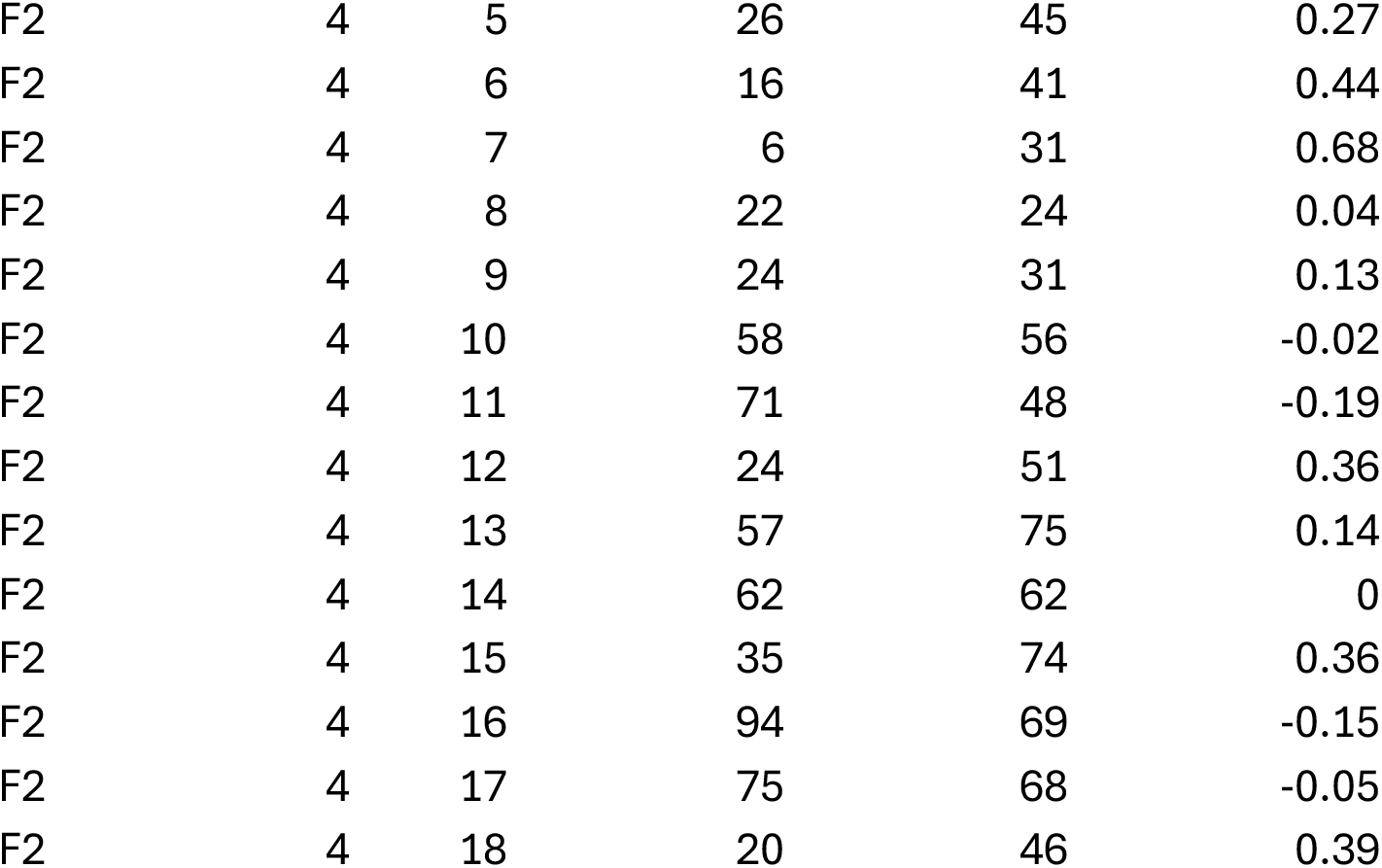
Raw data used in this study.

**Table S3.** Reagents and suppliers used in this study.

| Reagent | Supplier | Catalog |
| --- | --- | --- |
| **NGM Agar ingredients** |  |  |
| Agar | Carolina | 842133 |
| Cholesterol | FischerSci | AAA1147018 |
| MgSO<sub>4</sub> | FischerSci | 18-604-121 |
| CaCl<sub>2</sub> | FischerSci | 18-601-250 |
| KH<sub>2</sub>PO<sub>4</sub> | FischerSci | 01-337-803 |
| K<sub>2</sub>HPO<sub>4</sub> | FischerSci | 01-337-798 |
| NaCl | FischerSci | BP358-212 |
| Peptone | FischerSci | BP1420-500 |
| **Liquid NGM Buffer** |  |  |
| MgSO<sub>4</sub> | FischerSci | 18-604-121 |
| CaCl<sub>2</sub> | FischerSci | 18-601-250 |
| KH<sub>2</sub>PO<sub>4</sub> | FischerSci | 01-337-803 |
| K<sub>2</sub>HPO<sub>4</sub> | FischerSci | 01-337-798 |
| NaCl | FischerSci | BP358-212 |
| **LB Broth** |  |  |
| Yeast Extract | FischerSci | BP1422-500 |
| Tryptone | FischerSci | BP1421-500 |
| H<sub>2</sub>O (purified) | Barnstead Genepure | 10-451-061 |
| **Paralytic** |  |  |
| NaN<sub>3</sub> | FischerSci | S0489100G |
| **Bleach solution** |  |  |
| NaOH | FischerSci | S318-100 |
| NaOCl (6%) | Total Home Bleach | N/A |

